# Fungal extracellular vesicles are involved in intraspecies intracellular communication

**DOI:** 10.1101/2021.06.03.447024

**Authors:** Tamires A. Bitencourt, Otavio Hatanaka, Andre M. Pessoni, Mateus S. Freitas, Gabriel Trentin, Patrick Santos, Antonio Rossi, Nilce M. Martinez-Rossi, Lysangela L. Alves, Arturo Casadevall, Marcio L. Rodrigues, Fausto Almeida

**Affiliations:** Department of Biochemistry and Immunology, Ribeirao Preto Medical School, University of São Paulo, Ribeirao Preto, SP, Brazil; Department of Genetics, Ribeirão Preto Medical School, University of São Paulo, Ribeirão Preto, São Paulo, Brazil; Gene Expression Regulation Laboratory, Carlos Chagas Institute, Fiocruz, Curitiba, PR, Brazil; Department of Molecular Microbiology and Immunology, Johns Hopkins Bloomberg School of Public Health, Baltimore, MD 21205, USA; Instituto de Microbiologia Paulo de Góes, Universidade Federal do Rio de Janeiro, Rio de Janeiro, Brazil

**Author notes:** Corresponding author, (FA).

**Keywords:** Fungal infections, cellular communication, extracellular vesicles, virulence, fungal biology

## Abstract

Fungal infections are associated with high mortality rates in humans. The risk of fungal diseases creates the urgent need to broaden the knowledge base regarding their pathophysiology. In this sense, the role of extracellular vesicles (EVs) has been described to convey biological information and participate in the fungal-host interaction process. EVs play many roles, including cellular physiology, responding to environmental cues, mediating a complex circuit of cellular communication in bidirectional crosstalk with other organisms, and the communication between fungal cells has been speculated. This study demonstrated the intra species uptake of EVs in fungi, including *Candida albicans, Aspergillus fumigatus*, and *Paracoccidioides brasiliensis*, and the effects triggered by EVs in fungal cells. In *C. albicans*, we evaluated the involvement of EVs in yeast to hyphae transition, whilst in *P. brasiliensis* and *A. fumigatus* the function of EVs as stress transducers was investigated. Both *P. brasiliensis* and *A. fumigatus* were exposed to an inhibitor of glycosylation or UV light, respectively. The results demonstrated the role of EVs in regulating the expression of target genes and phenotype features. The EVs treatment induced cellular proliferation and boosted the transition yeast to hyphal transition in *C. albicans*, while they enhanced stress signals in *A. fumigatus* and *P. brasiliensis*, establishing a role for EVs in fungal intra species communication. Thus, fungal EVs regulate the virulence and adaptive traits in fungal interaction systems as potent message effectors, and understanding their effects and mechanism(s) of action could be exploited in antifungal therapies.

**Author Summary:** Extracellular vesicles (EVs) play an important role in export systems. They act as vehicles for the transference of complex cargoes with broad biological functions, such as proteins, carbohydrates, pigments, nucleic acids, and lipids. EVs can contribute to fungal infection outcomes. The EV content exerts immunomodulatory functions during fungus-host interactions. Furthermore, the participation of EVs in communication between fungal cells has been speculated. This study investigated the capacity of EVs to mediate intra-species in three genera of human pathogenic fungi and established a regulatory function of EVs. We also assessed the features of this regulation by analyzing the cellular morphological aspects of fungi after stimulation with EVs. Our data suggest fungal EVs can function as potent signal mediators that mediate virulence and adaptive responses.

## Introduction

Fungal infections are responsible for over 1.6 million deaths per year. It is estimated that more than a billion cases of severe fungal diseases affect the world population. Despite these numbers it is likely that they represent an underestimate of the fungal diseases that ail humans [1, 2]. The diseases caused by *Aspergillus spp., Candida spp*., and the endemic agent of mycoses such as *Paracoccidioides* species are among the deadliest mycoses [1, 2].

In recent years, extracellular vesicles (EVs) have been studied in cell-walled microorganisms. In fungi, they were first described in 2007 in *Cryptococcus neoformans* [3]. So far, EVs have been characterized in approximately 20 fungal species, including yeasts forms of *H. capsulatum, Sporothrix schenckii, C. parapsilosis, Saccharomyces cerevisiae, Malassezia sympodialis, P. brasiliensis, C. albicans, Pichia fermentans, C. gattii*, *S. brasiliensis, P. lutzii*, and *Exophiala dermatitidis* [4–13] and filamentous fungi such as *Alternaria infectoria, Trichophyton interdigitale*, *Rhizopus delemar, Fusarium oxysporum f. sp. Vasinfectum, Trichoderma reesei, Aspergillus fumigatus*, and *Aspergillus flavus [14–20]*.

EVs function as vehicles carrying complex cargoes with diverse biological functions, including proteins, carbohydrates, pigments, nucleic acids, and lipids. EVs can contribute to fungal infection outcomes [21]. EVs likely have roles in bi-directional communication, raising the possibility of communication between fungal cells [22]. Previous reports have demonstrated the participation of fungal EVs in biofilm formation [10], stimulation of cytokine production [6, 9, 11, 15, 20, 23], and favoring pathogen infection [4, 24]. Bidirectional communication mediated by fungal EVs has been demonstrated in the interaction of fungal cells with plants [25, 26] and in communication with mammalian cells [27, 28]. However, Intra fungal species communication mediated by EVs has not been reported so far.

The possibility of EV-mediated virulence transfer and/or antifungal resistance between strains have gathered attention. In the *Cryptoccocus* model, previous studies demonstrated that the Vancouver Island outbreak strain of *C. deuterogatti*, namely R265, can transfer its ability to proliferate within the host macrophages to an avirulent strain, which was attributed as an EV-regulated process [11, 29]. Another report showed that the supernatant from a highly virulent strain of *C. neoformans* culture stimulated the pathogenic potential of a less virulent isolate, an effect also attributed to EVs [30]. Recently, a study advocated the role of EVs from *C. albicans* as messaging compartments involved in growth, morphogenesis, and biofilm production [31]. These studies postulated a role for fungal EVs in intra-species communication; however, at the molecular level, many aspects related to the mechanisms switched on by EVs remain to be investigated. Herein, we sought to analyze the fungal cellular communication mediated by EVs using the fungal pathogens *P. brasiliensis, A. fumigatus*, and *C. albicans* using multiple approaches. Our data demonstrate that fungal EVs mediate cellular communication by regulating the expression of target genes and by controlling cellular proliferation.

## Results

### Intercellular transfer of fungal EVs

Because EVs can transport multiple molecules that play essential roles in fungal biology [22, 28, 32], we examined whether fungal EVs would be transferred intercellularly from cells of the same species. Using [1^−14^C] palmitic acid metabolic labeling, radiolabeled fungal EVs were produced as previously described [33, 34]. Radioactive assays confirmed that fungal EVs produced by *P. brasiliensis* (Fig 1A), *A. fumigatus* (Fig 1B), and *C. albicans* (Fig 1C) could be transferred from cell to cell within the same species. We tracked the radioactivity added to the fungal cell cultures during the time course after 0 h (control), 1 h, 6 h, 12 h, and 24 h from the addition of radiolabeled EVs to the cultures (Fig 1). The fungal cells were pelleted, washed with PBS, and pulse-chase measurements were performed. We verified that purified radiolabeled EVs were taken up by fungal cells as the radioactive signal increased over time.

**Fig 1.**
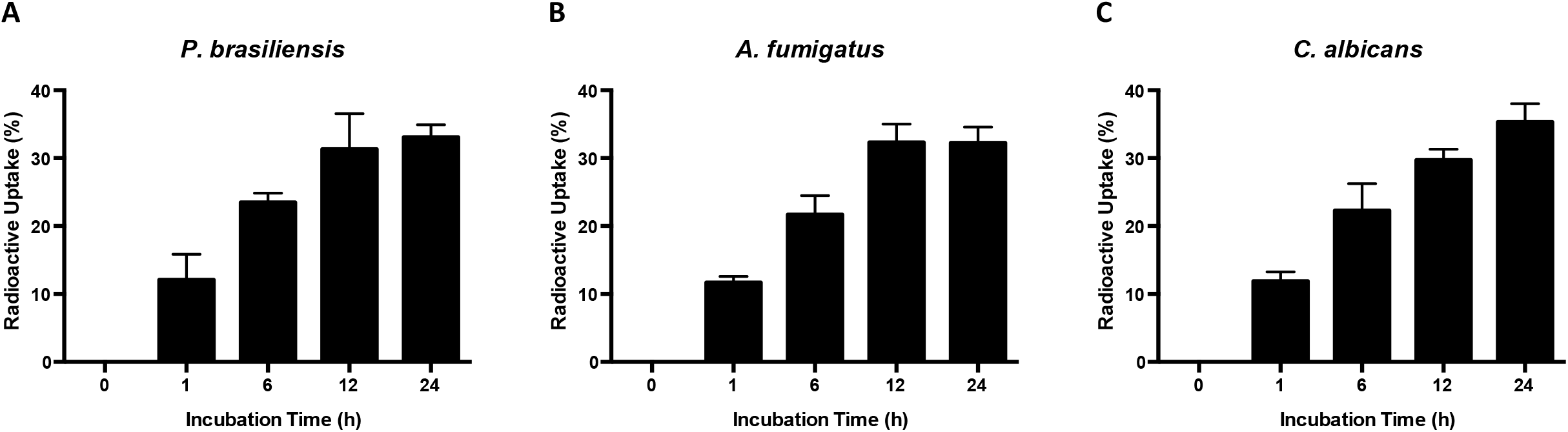
Evaluation of Extracellular Vesicles (EVs) uptake in different fungal species. The absorption of radioactive VEs was evaluated after 0h, 1h, 6h, 12h and 24h in the following yeast cells: *P. brasiliensis* (A), *A. fumigatus* (B) and *C. albicans* (C).

### *P. brasiliensis* EVs as cellular communicator during Endoplasmic Reticulum (ER) stress

We previously demonstrated that the genes PbHACA and PbIRE1 showed increased expression during tunicamycin (TM) treatment [35], suggesting that they are involved in the ER stress response of *P. brasiliensis*. Thus, we purified EVs from *P. brasiliensis* treated with TM (TM EVs) and added these EVs to *P. brasiliensis* yeast cells that did not receive TM treatment (Fig 2A). We observed that TM EVs increased PbHACA and PbIRE1 expression significantly (red bars, Figs 2B and 2C, respectively). On the other hand, fungal EVs obtained from *P. brasiliensis* yeast cells that were not treated with TM (Pb EVs) added to *P. brasiliensis* yeast cells without TM treatment did not alter PbIRE1 and PbHACA expression (blue bars, Figs 2B and 2C, respectively). PBS was used as a negative control (white bar), while TM was the experimental group (black bars). These results strongly suggested that *P. brasiliensis* EVs could participate in intercellular communication during ER stress, possibly promoting fungal adaptive responses.

**Fig 2.**
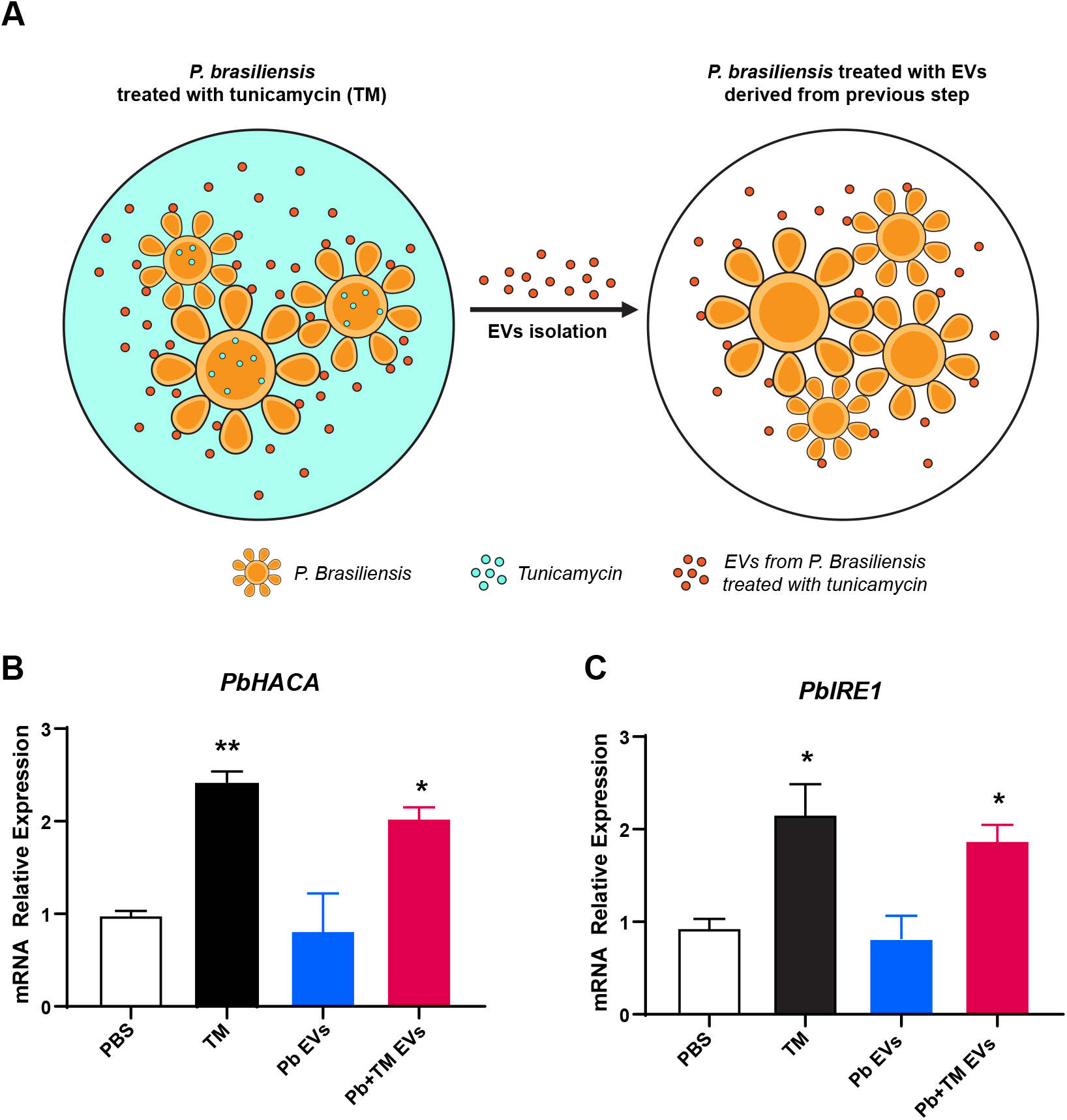
EVs from *P. brasiliensis* as cellular communicator during ER stress. Schematic representation of EVs obtained after tunicamycin exposure (A). The relative expression of UPR belonging genes *IRE1* (B) and *HACA* (C) were determined after EVs uptake. Significantly different values are indicated by asterisks as determined using ANOVA followed by Tukey’s post hoc test (P<0.05). The calibrator was PBS condition and the positive control was tunicamycin (TM).

### EVs from *A. fumigatus* act as stress message effectors

To verify whether the observations made with *P. brasiliensis* applied to other fungi, we isolated EVs from ultraviolet (UV) irradiated cultures and from non-irradiated *A. fumigatus* cells (control). The EVs obtained from UV-treated cells were named UV EV, whereas the EVs obtained from regular cultures were named Control EV. Our data showed an overall population size in the range of 100 to 200 nm and a minor population of EVs with sizes varying from 320 to 394 nm in regular cultures or a range of 240 to 614 nm in cultures that underwent UV irradiation (S1 Fig).

EVs obtained from both *A. fumigatus* cultures after UV irradiation (UV EV) or without UV irradiation (Control EV) (Fig 3A) caused a prominent decrease in colony formation in *A. fumigatus* (Fig 3B). Approximately 40 percent of colony reduction was achieved after EVs uptake (Fig 3B). Thereafter, the gene expression analysis reinforced a possible role of these EVs as stress message effectors, showing the upregulation of the *mpkC* gene in both EVs uptake conditions. The *mpkC* gene plays a role in adaptive responses to different stress agents such as osmotic stress, oxidative stress, heat shock, cell wall damage and also acts in cellular division regulation, as demonstrated in *A. fumigatus* [36, 37]. The cultures that received UV EV showed higher induction of the *mpkC* gene (Fig 3C). In addition, UV EV also caused a significant increase in the *akuA* transcript levels (Fig 3D), supporting a role of EVs in cellular communication events. In addition, the presence of EVs caused a subtle downregulation of *the nimA* gene, which may reflect another level of cell cycle regulation (Fig 3E).

**Fig 3.**
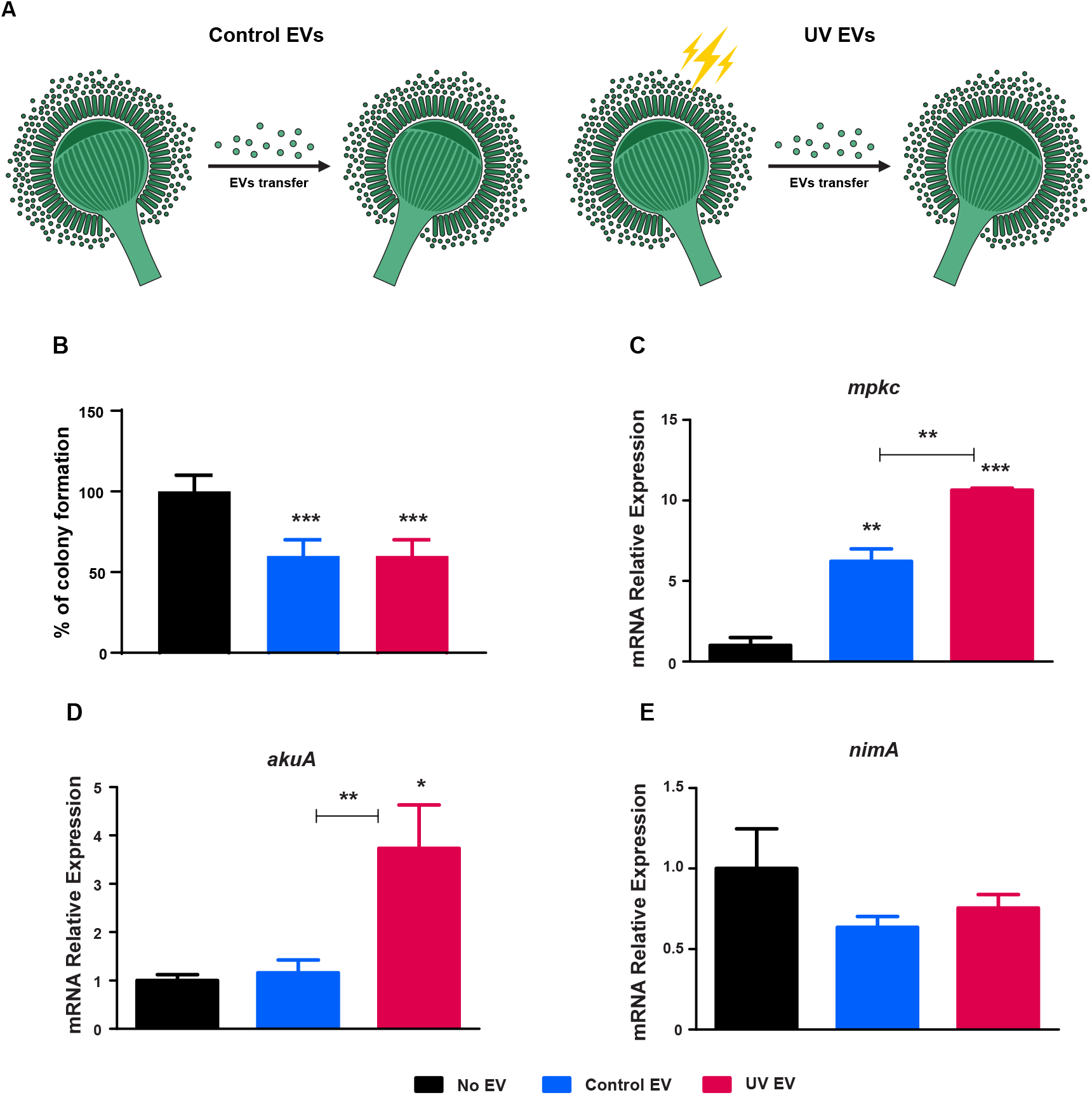
EVs from *A. fumigatus* as stress message effectors. Schematic representation of EVs obtained from regular cultures, without UV exposed (Control EV), and EVs obtained from UV exposed cultures (UV EV) (A). The thunder diagram represents UV light exposure. The colony formation was counted and a percent of reduction was determined after EVs uptake (B). RT-qPCR for *A. fumigatus* genes *mpkC* (C), *akuA (D)* and *nimA* (E). Relative expression was assessed using NO EV uptake condition as reference sample, and 18S and *βtub* as reference of normalization. Significantly different values are indicated by asterisks as determined using ANOVA followed by Tukey’s post hoc test (*P < 0.05, **P<0.01, ***P<0.001).

### EVs induce hyphae formation and regulate the cell cycle in *C. albicans*

Next, we investigated whether EVs obtained from yeast culture and a yeast to hyphae culture in the *C. albicans* model would mediate communication signals that affect its morphological transition. Therefore, we isolated EVs from *C. albicans* from culture conditions of hyphae formation and from yeast cultures. We analyzed the size and distribution of EVs in both cultures (S1 Fig). EVs obtained after *C. albicans* grown on cultures in YPD at pH 6.3 at 30 °C were named as Control EV, whereas EVs obtained from *C. albicans* grown on cultures in YPD at pH 7.4 at 37 °C were named Trans EV. Our results showed a heterogeneous EV distribution profile within these conditions, and the majority of EVs obtained from yeast to hyphae transition cultures (Trans EV) had a range size from 99 to 182 nm and minor populations with sizes varying from 23 to 85 nm and 232 to 440 nm. For EVs isolated from *C. albicans* yeast cultures (Control EV), the majority of EVs had a range size from 107 to 154 nm, and a minor population in the range of 211 to 440 nm. Previous studies have demonstrated the heterogeneity of EVs from *Candida spp*., showing a distribution profile with population sizes ranging from 60 to 280 nm, and in *C. albicans*, the sizes varied from 100 nm to up 600 nm [5, 38, 39].

We also evaluated the gene expression profiles of selected genes (*hwp1, sap5, cht2*, and *sec24*) during *C. albicans* growth in yeast to hyphae transition cultures in comparison to yeast cultures after different incubation times, such as 0.5, 1, 2, and 4 h (S2 Fig). These genes were previously modulated in microarray data of *C. albicans* grown as hyphae with serum at 37 °C, and *sap5* (“secreted aspartyl protease”) and *hwp1* (“cell wall protein hyphal wall protein 1”) were highly upregulated [40]. We identified a time-dependent modulation for the genes *cht2* and *sec24*, with a significant downregulation after 4 h of growth. A prominent upregulation was verified for the *hwp1* gene at all time points evaluated. The *sap5* gene was upregulated at all times analyzed. Taking advantage of this information, we compared gene expression profiles after EVs uptake in *C. albicans*. Thus, the *C. albicans* cells were incubated with Trans EV or Control EV (Fig 4 A). Thereafter, the cells were recovered through centrifugation and resuspended in a yeast to hyphae medium. By analyzing the morphological features of *C. albicans* in yeast-to-hyphae condition assessed throughout the incubation time points, we identified the occurrence of the three main morphologies presented by *C. albicans*, yeast, pseudo-hyphae, and hyphae. Our results also showed a possible increase in cellular proliferation after EVs uptake compared to no EV uptake cultures. The XTT assay reinforced the involvement of EVs in cell cycle regulation. (Fig 4B). Moreover, the culture that underwent Trans-EV uptake appeared to present changes in yeast development, in which a particular cellular clumping was detected even at earlier time points (Fig 4C).

**Fig 4.**
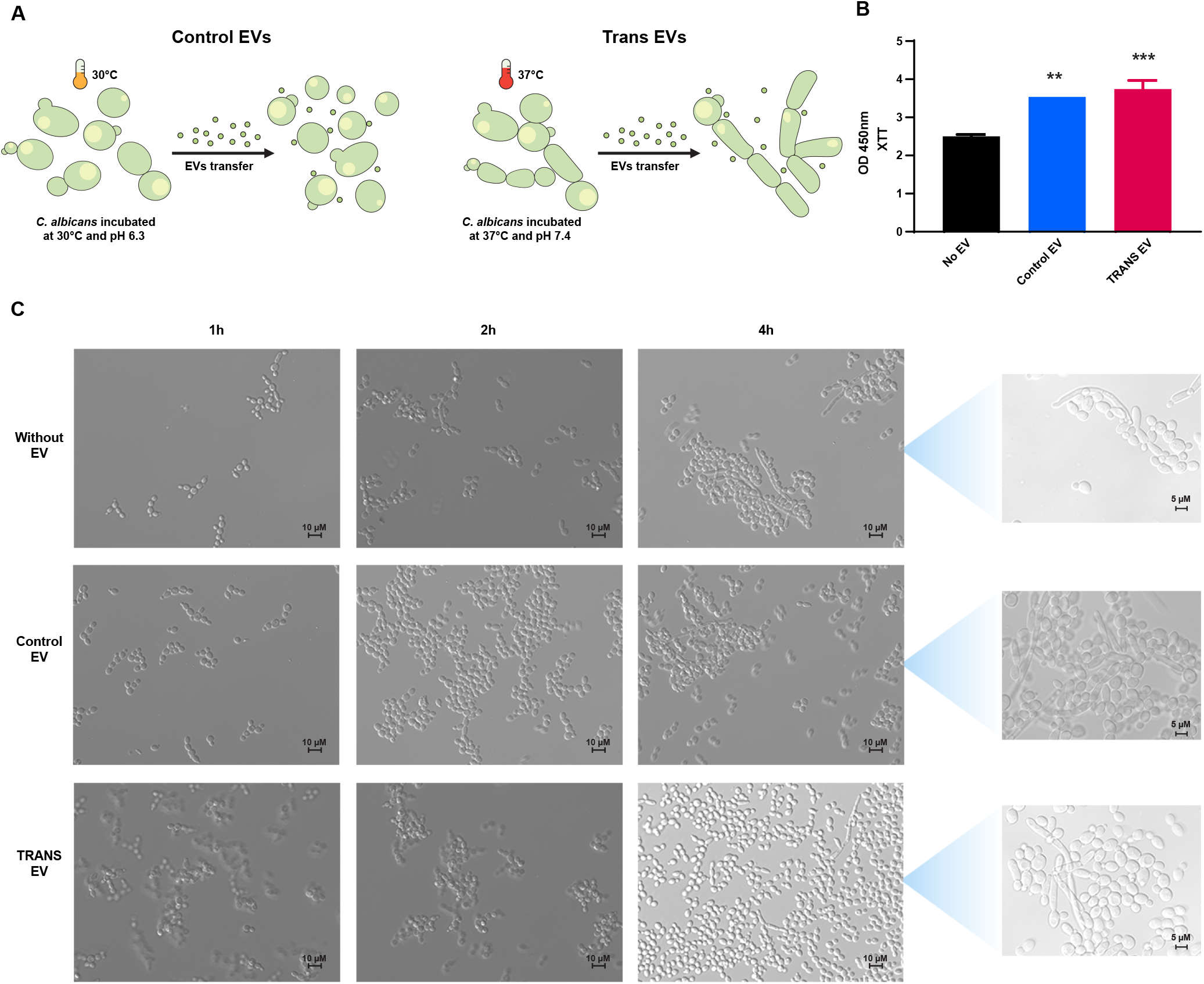
EVs heighten the cellular proliferation in *C. albicans*. Schematic representation of EVs obtained from yeast cultures (Control EV) and yeast to hyphae cultures (Trans EV) (A). XTT reduction assay to assess cellular proliferation after EVs uptake during 2h of growth on transition cultures (B). Morphological appearance addressed by microscope images of *C. albicans* cells grown on YPD pH 7.4 at 37°C after EVs uptake (C).

EV uptake caused a rapid response toward hypha-inducing stimuli, with a more significant upregulation of the *hwp1* gene at an earlier incubation time point (1h) followed by a decay in its transcript levels at a later time point (4h), which was observed for both uptake conditions, Control EV and Trans EV (Figs 5 A and 5 E). The transition panel (S2 Fig) suggested a fluctuation in the *hwp1* modulation, with a higher increase in the transcript levels after 2h, showing an approximately 50-fold difference. In addition, the EVs from transition cultures (Trans EV) promoted repression of the *sec24* gene after 1 and 4 h (Figs 5B and 5F). The transition panel demonstrated its downregulation, particularly in delayed incubation time points, in which hyphae were more commonly found. In addition, the *cht2* gene was repressed after EVs uptake from transition cultures after 1 h, and no change in its transcription levels was observed after 4h (Figs 5C and 5G). A previous study showed that the downregulation of this *cht2* gene is a remarkable feature in hyphae [41]. Therefore, our data suggest a boost in hyphae induction after EVs stimulation. Curiously, *sap5*, which is mainly upregulated in cells with hyphal morphology, presented a decrease in the transcription levels in EVs uptake conditions (Figs 5D and 5H).

**Fig 5.**
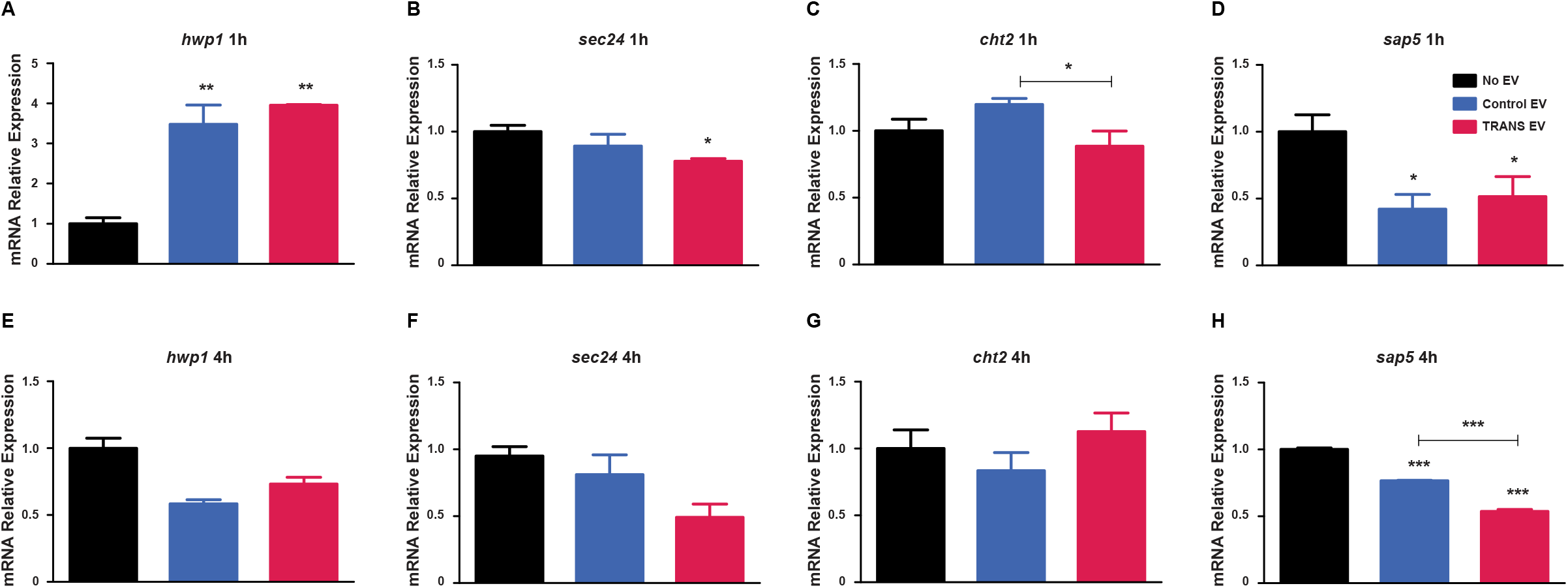
EVs prompted a boost in yeast to hyphae transition gene expression response. RT-qPCR evaluated a set of *C. albican*s genes after EVs uptake during 1h and 4h of growth on transition cultures. *hwp1* (A and E)*, sec24* (B and F)*, cht2* (C and G) and *sap5* (D and H). The relative expression was assessed using NO EV uptake condition as the reference sample after normalization with the *rpp2b* and *tdh3* genes. Significantly different values are indicated by asterisks, determined using ANOVA followed by Tukey’s post hoc test (P< 0.05, ^**^P< 0.01, ^***^P<0.001).

## Discussion

Intercellular communication occurs through contact with cells or by the secretion of molecules such as quorum sensing (QS) effectors. A recent study demonstrated that EVs enriched with Fks1 and Chs3 from *Saccharomyces cerevisiae* rescued the yeast cells from cell wall disturbing [42], which suggested the possibility that EVs function in fungal intra-species communication.

Herein, we evaluated whether EVs are capable of mediating intra-species communication in fungi by applying three approaches in three different fungal pathogens. We used an uptake time of 1 h, which was sufficient to allow the incorporation of radioactive EVs in different fungal cells, as aforementioned (Fig. 1). It is expected that subtle differences in EVs uptake could be responsible for altering and regulating gene expression because fungi have the refined ability to sense any change in the environment and adjust their metabolism in response to these changes.

Our study demonstrated a role for EVs as stress message effectors in the fungal agents *P. brasiliensis and A. fumigatus*. In the *P. brasiliensis* model, the genes belonging to the UPR pathway were studied after cultures underwent EVs uptake (EVs obtained from previous tunicamycin-exposed cultures) in comparison to cells directly exposed to tunicamycin. The UPR pathway is switched on to re-establish Endoplasmic Reticulum homeostasis by heightening the folding capability and controlling misfolded protein disposal [43]. In fungi, this pathway comprises the ER-transmembrane sensor Ire1/IreA (Ser/Thr kinase) and a transcription factor Hac1/HacA [44]. Our experiments demonstrated the upregulation of both genes in EVs uptake cultures, suggesting that EVs function in fungal communication.

In the *A. fumigatus* model, EVs obtained from a regular or an irradiated culture triggered a stress response in fresh *A. fumigatus* cultures, as evidenced by colony reduction formation and upregulation of a stress response gene, the *mpkC* gene. We hypothesize that fungitoxic compounds in EVs from *A. fumigatus* may be responsible for these effects. Previously, the production of antifungal compounds by filamentous fungi such as *A. fumigatus* was reported [45]. However, the characterization and understanding of the processes mediated by EVs in *A. fumigatus* have been poorly explored. Recently, a study showed the production and cargo description of EVs from *A. fumigatus*, in which approximately 60 proteins were identified with potential roles in immunomodulation and pathogenicity [19].

In this study, the uptake of EVs from cultures exposed to UV light prompted the most pronounced upregulation of the *mpkC* gene. It also promoted a remarkable increase in transcript levels of the *akuA* gene. *akuA* is involved in DNA repair [46]. UV exposure causes DNA damage through mutation and pyrimidine dimers and induces oxidative stress status [47, 48]. A previous study demonstrated the involvement of *akuA* in cellular protection against UV radiation exposure, showing that *akuA* deletion was responsible for a significant reduction in survival rates of the mutant strain compared to *A. fumigtaus* wild type [49]. Moreover, we revealed that the cultures that underwent EVs uptake from *A. fumigatus* showed a trend of downregulating an encoding gene of cyclin-dependent kinase, NimA, involved in cell cycle transition [50].

In *C. albicans*, we assessed the potential of EVs to favor hyphal formation. It is widely accepted that dimorphism in *C. albicans* is important for virulence [51]. The yeast-to-hyphae transition is a tightly regulated process of sensing and responding to environmental cues [52]. Hyphae formation is mainly associated with invasion and adaptive responses during fungus-host interactions [53]. Recently, single-cell RNA-seq data showed the dynamic regulation of transition genes during macrophage infection with *C. albicans* and also highlighted the occurrence of bimodal modulation in genes related to hyphae formation and cell wall remodeling concomitantly with differences in immunomodulation responses [54].

In our data, we detected a new response characterized by cell clumping after Trans EV uptake, in which the EVs were obtained from *C. albicans* hyphae-inducing cultures. Cell clumping changes probably reflect changes in surface hydrophobicity. This feature may result from cell surface changes activated in the regulation of hyphal expansion [55, 56]. Furthermore, the gene modulation profile assessed after EVs uptake also showed a boost in yeast to hyphae transition response, mainly prompted after Trans EV uptake, which is reinforced by the presence of long hyphae in this treatment (S3 Fig). Hyphal growth is associated with a hyphal-specific transcriptional program that paves the way towards the expression of genes related to virulence traits such as adhesion (ALS3 and HWP1), invasion (ALS3), oxidative stress response (SOD5), proteolytic activity (SAPs), and others [53]. In a previous high-throughput study, a set of genes showed differential modulation after transition stimuli, including the genes *sap5, hwp1, sec24, cht2* [40]. Furthermore, hyphal development relies on a complex network of transduction signals that respond to environmental cues and activate the molecular processes involved in driving the high polarization growth of hyphal forms [57, 58].

Aspects of cell cycle organization are also unique in hyphae compared to pseudo-hyphae and yeasts. Hyphal forms showed different branching patterns, regulation of the cell cycle, and polarization [58]. In addition, a phenomenon known as QS governs responses related to hyphal formation and control of cell density [59, 60]. Many QS molecules might also be found within fungal EVs; however, it is also possible to speculate a cross-regulation between small RNA content in EVs and QS phenomena in fungi. A recent study showed the involvement of QS molecules such as farnesol derivatives and medium-chain fatty acids in EVs obtained from yeast culture as potential regulators of cellular proliferation and yeast-to-hyphae development [31]. The EVs from *C. albicans* reduced the hyphae and biofilm formation, and the impact of *C. albicans* EVs on invasive hyphae development and virulence, as shown in *Galleria mellonella* model [31]. Moreover, that study was conducted exploring EVs obtained from yeast cultures, and it is reasonable that the cargo and the message within this EV contribute to cells remain in this form.

The current knowledge regarding the differences in cell cycle regulation in hyphal development associated with our data prompted the investigation of cellular proliferation after EVs uptake. We demonstrated that the conditions that underwent EVs uptake presented an increase in mitochondrial activity, suggesting a potential function of EVs in heightening cellular metabolism and/or proliferation. EVs were previously shown to enhance cellular proliferation in *C. albicans* [31] as well as in *C. neoformans* within macrophages. The high activation of the *C. neoformans* proliferation rate was named the “division of labor” mechanism [11]. It was ascribed as an important virulence trait transferred from an outbreak strain to a non-outbreak strain [11].

Collectively, our data pave new avenues for EVs functions in fungal intra-species communication, supporting adaptive responses and virulence mechanisms. Although we recognize the need to develop our knowledge about the complex cellular communication circuit switched on by EVs, and regardless of how it will be addressed in future, this study sheds new light on the role of fungal EVs in intra-species cellular communication.

## Methods

### Fungal strains and growth conditions

*Paracoccidioides brasiliensis* strain 18 (Pb18) was cultivated as previously described [61]. The Pb18 yeast form was maintained in Fava Netto semi-solid medium and incubated at 36°C. Yeast growth was performed by inoculating cultures in liquid YPD medium (2% peptone, 1%yeast extract, and 2% glucose) at 36 °C on a rotary shaker at 100 rpm for 72 h. Tunicamycin (TM) was used to induce endoplasmic reticulum (ER) stress. TM (15 μg/ml) was prepared in 20 mM of NaOH, and it was added to yeast cultures in liquid YPD for 5 days at 36 °C on a rotary shaker at100 rpm to obtain EVs derived from ER-stressed fungi. The same fungal growth was performed without TM to obtain Pb18 control EVs.

The *Aspergillus fumigatus* strain CEA17 used in this study was grown in a complete agar malt medium [YAG medium supplemented with malt extract: 2% (w/v) glucose, 0.2% (w/v) yeast extract, 2% (w/v) malt extract, 2% (w/v) agar, and 0.1 % (v/v) trace elements] for 7 days at 37 °C. Conidia suspensions were obtained from 7-day-old plates with sterilized PBS, recovered by centrifugation, and filtered through sterile Miracloth (Millipore, Billerica, MA, USA). Conidia concentration was determined using a Neubauer chamber. Approximately 1 × 10^3^ conidia were directly inoculated into a complete YAG medium or the conidia suspension was first exposed to UV light for 60 s and then inoculated into a complete YAG medium. The cultures were incubated for two days at 37 °C. These plates were used for the following experiments: EVs isolation and communication assay.

*Candida albicans* strain ATTC-64548 was grown at 30 °C in Sabouraud medium (Oxoid, Basingstoke, UK) for 72h. One fresh colony was inoculated into 10 ml of yeast extract-peptone-dextrose medium [YPD: 1% (w/v) yeast extract, 2% (w/v) peptone, 2% (w/v) dextrose, pH 6.3], and cultured overnight at 30 °C with shaking (150rpm). Then, the overnight cultures of *C. albicans* were diluted to an OD_600_ range of 0.100-0.130 with YPD medium (pH 6.3) or YPD medium (pH 7.4), and grown at 30 °C or 37 °C, respectively. The cultures grown on YPD at pH 6.3 represent the control cultures, in which no stimuli for hyphae differentiation were offered, whereas cultures grown on YPD at pH 7.4 represent the transition cultures, in which a stimulus conferred by pH (7.4) and temperature (37°C) was offered to prompt hyphae differentiation. After that, the cultures were used to perform the EVs isolation experiment, the gene expression profile of yeast to hyphae, and the communication assay.

### EVs isolation

EVs from Pb18 yeast cultivated in the presence or absence of TM were isolated as previously described [62]. Yeast cultures of Pb18 growth in YPD medium were depleted from cell pellets by serial centrifugation at 5,000 × g for 15 min and 15,000 × g for 30 min at 4 °C. The supernatants, containing EVs, were concentrated and filtered using Amicon^®^ systems (Millipore, Billerica, MA, USA) with a 100-kDa cutoff membrane. The concentrated material was centrifuged again at 15,000 × g for 30 min at 4 °C, and the supernatants were centrifuged at 100,000 × g for 1 h at 4 °C to collect vesicles. The EVs pellets were resuspended in PBS for NTA analysis and experiments. EVs obtained from TM-treated Pb18 were termed TM EVs, whereas EVs obtained from non-treated cultures were termed Pb EVs.

Regarding the isolation of *A. fumigatus* EVs, the procedures were performed as previously described [19, 39], with slight modifications. 2-days-old plates obtained for a regular culture or a UV-exposed culture were used to isolate EVs. UV light exposure cultures were obtained after conidial irradiation. Approximately 10^4^ conidia/ml were irradiated with UV germicidal light (G1578 UV lamp) at 16 cm distance for 60 s with constant shaking, and then 100 μl of this irradiated suspension were plated on YAG medium, yielding about 10% of CFUs in comparison to non-irradiated cultures. The cells were washed twice with 3 ml of sterile PBS, recovered from the dish plates with inoculation loops, and then transferred to centrifuge tubes. Thereafter, the cell suspension was filtered through sterile Miracloth (Millipore, Billerica, MA, USA), and then a sequential centrifugation and supernatant concentration in the Amicon system was performed. The EVs obtained from UV light exposure were named UV EV, whereas the EVs obtained from regular cultures were called Control EV.

Isolation of *C. albicans* EVs was performed as described previously, with slight modifications [9, 42]. The concentration of the *C. albicans* cultures was adjusted as mentioned above, and the preinocula were added into 300 ml of YPD (pH 6.3) or YPD (pH 7.4), and incubated for 4h at 30 °C or 37 °C, with shaking (100 rpm). For EVs isolation, the cells and debris were removed by sequential centrifugation at 4000 × g for 15 min and 15,000 × g for 15 min. Supernatants were concentrated using an Amicon ultra-concentration system (cutoff 100 kDa, Millipore, Billerica, MA, USA). The resulting concentrated supernatant was ultracentrifuged at 100,000 × g for 1 h at 4 °C. Pellets were collected and resuspended in ultra-pure water (Sigma-Aldrich, St. Louis, MO, USA) supplemented with protease inhibitor cocktail 10X (Sigma) (0.2% v/v), and stored at −80 °C. Taking into account the culture conditions used to promote yeast-to-hyphae transition (YPD medium at pH 7.4, and incubated at 37 °C), an additional step was included, in which the cultures were first filtered using sterile Miracloth (Millipore, Billerica, MA, USA), and then subjected to differential centrifugation. EVs obtained after *C. albicans* grown on YPD at pH 6.3 at 30 °C were named as Control EV, while EVs obtained from *C. albicans* grown on YPD at pH 7.4 and incubated at 37 °C were named Trans EV.

### Nanoparticle-Tracking Analysis (NTA)

To determine the size distribution and quantification of EVs isolated from *C. albicans* cultures in two different stages, a yeast culture condition (Control EV) and a yeast-to-hyphae culture condition (Trans EV), NTA analyses were performed. We also performed the NTA analysis to measure and characterize the size distribution of EVs isolated from regular and UV-exposed *A. fumigatus* cultures. We obtained the EV profiles of *P. brasiliensis* EVs, as previously described [34].NTA analysis was performed using a Nanosight appliance NS300 (Malvern Instruments, Malvern, UK) with NTA 3.0. The NTA analysis is an optical dispersion technique employed to measure the size distribution of particles in a solution at the nanometer scale [63].

### Determination of EVs Uptake by yeast cells

To obtain radiolabeled EVs, each one of the fungal species (*P. brasiliensis*,*A. fumigatus*, and *C. albicans*) was pulsed with [1-^14^C] palmitic acid for 72 h before EVs isolation [64]. The culture medium used for each species, and EVs isolation process were described above. Radiolabeled EVs from the same species (10^9^/ml) were added to the corresponding yeast phase fungi (10^7^/ml) and incubated for 0, 1, 6, 12, and 24h at 37 °C in 9.5% CO_2_. After incubation, fungal cells were washed in PBS and lysed in 200μl of 25 mM deoxycholate, and the resultant material was collected for scintillation counting.

### Gene expression profile of *C. albicans* in yeast to hyphal transition stage

The *C. albicans* inoculum was prepared and its concentration was adjusted as previously described. It was added to 5 ml of YPD (pH 6.3) and incubated for 0.5, 2, and 4 h at 30 °C (yeast culture condition). Thereafter, the cultures were centrifuged at 4000 × g for 10 min at 4 °C. The resulting pellet was stored at - 80 °C until RNA extraction. Alike, the *C. albicans* inoculum was also added to 5 ml of YPD (pH 7.4) and incubated for 0.5, 2, and 4 h at 37 °C (hyphae-yeast culture condition). The pellet obtained from these cultures was stored at - 80 °C until RNA extraction. We used the resulting material to create a panel that reflects the expression profiles of *hwp1, sec24, sap5*, and *cht2* genes during *the C. albicans* transition stage.

### Communication assay mediated by EVs

#### Effect of EV on *P. brasiliensis* endoplasmic reticulum stress

The *P. brasiliensis* yeast cells were treated with TM, as previously described [35]. The EVs from TM-treated and untreated yeast cultures were isolated and the uptake of EVs (4×10^9^/ml) with *P. brasiliensis* yeast cells (10^5^/ml) was performed for 2 h at 37 °C in BHI medium (Sigma). RNA isolation, cDNA synthesis, and qPCR assays were performed as described previously [35].

#### Effects of EV on *A. fumigatus* UV stress

This assay was performed with 5 ml of *A. fumigatus* conidia suspension (10^4^ conidia/mL). The conidia suspension was incubated with 5 × 10^8^ EVs/ml obtained from *A. fumigatus* regular cultures or UV light-exposed cultures. The uptake was performed for 1 h at 37 °C in PBS with shaking (100 rpm). Therefore, the fungal cells were recovered by centrifugation at 4000 × g for 10 min at room temperature and the cultures were resuspended in 5 ml of PBS. Subsequently, approximately 100 μl of conidia were plated on the YAG medium. After incubation at 37 °C in the dark for 2 days, the CFUs were counted. CFUs from conidia without EV uptake were counted as controls. The colonies were stored at - 80 °C until RNA extraction.

#### Effects of EV on *C. albicans* dimorphism

About 5 × 10^8^ EVs obtained from *C. albicans* yeast cultures or *C. albicans* yeast-to-hyphae transition cultures were added into a fresh culture of *C. albicans* with the cell density adjusted to an OD_600_ of 0.100-0.130, as previously mentioned, in 5 ml of YPD (pH 6.3 for 1h at 30 °C), with shaking (100 rpm). The yeast cells were then recovered by centrifugation. The pellets were then resuspended in 5 ml of YPD pH 6.3 and incubated for 1h and 4h at 30 °C with shaking (100 rpm). Alike, the pellets were also resuspended in 5 ml of YPD (pH 7.4) and incubated for 1h and 4h at 37 °C under shaking (100rpm). After incubation, the cultures were centrifuged at 4000 × g for 10 min at 4 °C, and the pellet was stored at −80 °C until RNA extraction. Similar conditions were employed to analyze cellular proliferation through the 2.3-bis (2-methoxy-4-nitro-5-sulfophenyl)-5-[carbonyl (phenylamino)]-2H-tetrazolium hydroxide (XTT) reduction assay. The XTT assay was performed as previously described [65] with slight modifications. After the uptake of EVs in *C. albicans* cultures, the yeast pellets were recovered through centrifugation at 4000 × g for 10 min at room temperature, and the pellets were resuspended in YPD (pH7.4) and incubated for 2h at 37°C. Then, the medium was removed, and the pellet was resuspended in XTT solution (1 mg/ml in PBS) and menadione (1mM in acetone), and incubated for 3h with gentle shaking. The activity of the yeast mitochondrial dehydrogenase reduces the tetrazolium salt XTT to formazan salts, which results in a colorimetric change that might correlate with cell viability. Colorimetric changes were measured using an ELISA microplate reader (MULTISKAN FC, Thermo Scientific) at 450 nm. Additionally, we followed the changes in yeast-to-hyphae transition in *C. albicans* cultures after EVs uptake by microscopy analysis. Images were obtained using Zeiss Observer Z.1 microscope with AxioVision SE64 software after 1h, 2h, and 4h time points. As controls, cultures that had not undergone the EVs uptake process were analyzed.

### RNA extraction and quantitative real time PCR (RT-qPCR)

The *C. albicans* cells were treated with lysis solution (20 mg/ml lysozyme, 0.7 M KCl, and 1 M MgSO4, pH 6.8) for 1 h with shaking (100 rpm). Next, the supernatant was removed by centrifugation at 1000 × *g* for 10 min, and total RNA was extracted using the Illustra RNAspin Mini RNA Isolation Kit (GE Healthcare), following the manufacturer’s instructions. The *A. fumigatus* mycelia were lysed by mechanical pulverization with a pestle and mortar in liquid nitrogen, and total RNA extraction was performed using Illustra RNAspin Mini RNA Isolation Kit (GE Healthcare). *P. brasiliensis* cells were also lysed with the aid of a small mortar and pestle in liquid nitrogen before total RNA was isolated using Trizol reagent (Life Technologies, Carlsbad, CA, USA), as described previously [35]. RNA concentration and quality were estimated using a nanophotometer (Implem). The RNA was pretreated with DNase (Sigma). Complementary DNA (cDNA) synthesis was performed using the High-Capacity cDNA Reverse Transcription kit (Applied Biosystems) following the manufacturer’s instructions. Quantitative RT-PCR was conducted as described previously [66]. The qPCR experiments were performed with SYBR Green Master Mix (Applied Biosystems) in the Step One Plus platform. Primer sequences were retrieved from the IDT DNA “primer quest” tool (www.idtdna.com/primerquest/Home/Index), and the oligonucleotide sequences are listed in Table 1. The algorithm used for gene expression analysis was the relative quantification 2^−ΔΔCt^ method [67], and graphs were generated using GraphPad Prism v.5 software (GraphPad).

**Table 1.**
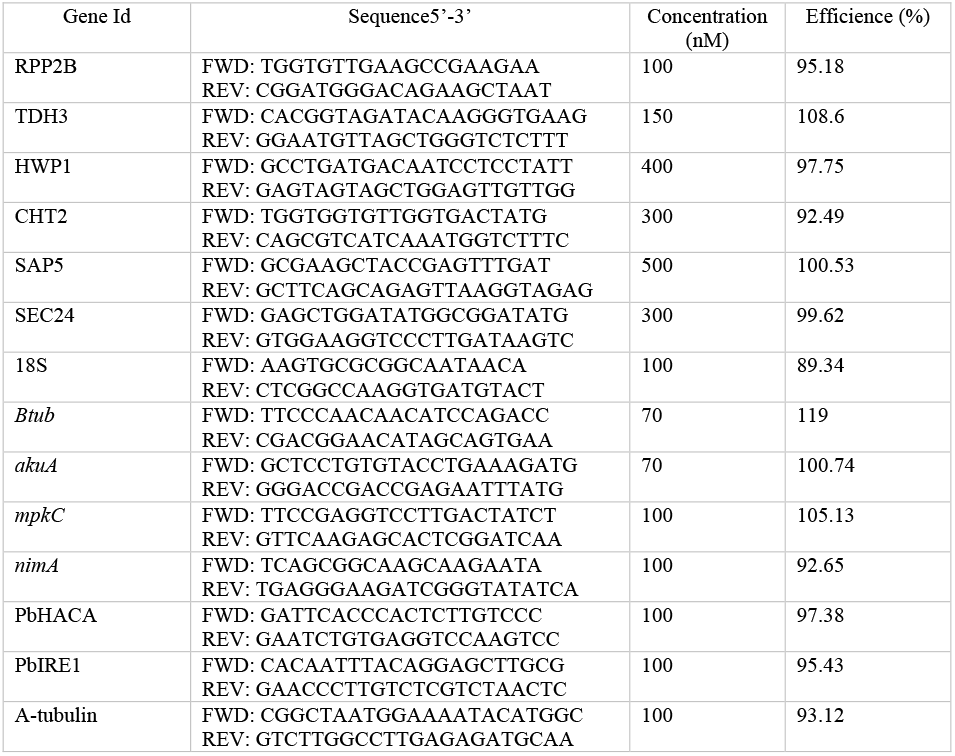
Set of primers used in RT-qPCR.

### Statistical analyses

The results are presented as mean values from independent experimental ±standard deviations. Significant differences were determined by one-way analysis of variance followed by Tukey’s post hoc tests or by unpaired t-test, using the GraphPad Prism 5 software.

## Author Contributions

FA, OH, AC, MR, and TB conceived the study. FA, OH, and TAB. performed the experimental design and laboratory experiments. AP, and MSF contributed with *A. fumigatus* assays. AP and PS performed the illustrative designs. GT assisted in the *C. albicans* EVs isolation assay. TB, NM, AR, AC, LA, MR and FA wrote the manuscript.

## ACKNOWLEDGMENTS

We thank Carlos Alberto Vieira for technical support.

## Supplementary Content

**S1 Fig.** Histograms from particle-size distribution of extracellular vesicles (EVs) from *C. albicans* and *A. fumigatus* for each assessed condition.

**S2 Fig.** Transition gene expression profile in *C. albicans* for selected genes such as *hwp1, sec24, cht2*, and *sap5* after 0.5, 2, and 4h. Control cultures were used as reference for the modulation of transition cultures. *The rpp2b* and *tdh3* genes were used as normalizer genes. Significantly different values are indicated by asterisks, as determined using an unpaired t-test (*P < 0.05).

**S3 Fig.** Morphological traits evaluated after EV treatment at 4 h in yeast to hyphae transition condition. The fungal forms were examined by optical microscopy. Control with No EV (A), treatment with Control EV (B) and treatment with Trans EV (C). Hyphae are shown by blue arrows and pseudo-hyphae are represented by black arrows.

